# Evolution of pollination-induced plasticity of flower longevity

**DOI:** 10.1101/2023.03.07.531535

**Authors:** Kuangyi Xu

## Abstract

In many plants, pollen deposition or removal can lead to early flower senescence, while flowers will live longer when pollination is excluded, and the strength of this pollination-induced plasticity of flower longevity varies across populations. This study builds models to investigate the evolution of the plasticity of flower longevity through group and individual selection under stochastic pollination environments. Generally, stronger plasticity evolves when female and male fitness accrual rates are higher and more volatile, and the evolution of plasticity is also strongly affected by daily maintenance cost of flowers relative to construction costs. Group selection only selects for plasticity in the female function (i.e., ovule fertilization) but not in the male function (i.e., pollen removal), and plasticity is optimal in maximizing fitness only when saved resources through plasticity greatly enhance individual fitness. Plasticity is more likely to evolve under individual selection, but under the same pollination environment, any combination of the plasticity in female and male functions that adds up to a certain overall strength of plasticity can be evolutionarily stable. Given the fact that plasticity in the male function is rare in hermaphroditic populations, the plasticity of flower longevity may evolve mainly through group selection instead of individual selection.

## Introduction

Flower longevity is an important factor for the reproductive success of plants, since it influences the number of pollinator visits (Ashman 1994, Primack 1985, van Doorn 1997, van Doorn and Kamdee 2014). Plant species exhibit great variations in flower longevity, with flowers of some species (e.g., cactus) persist only for several hours while flowers of other species (e.g., many orchids) can keep open for several months. Although longer-lived flowers tend to increase the opportunity of pollen deposition and removal, it costs resources (e.g., water) for the maintenance of flowers, which limits the number of flowers constructed (Spigler and Woodard 2019). Based on the resource allocation theory, evolutionary models of flower longevity show that larger flower longevity is selected under lower rates of pollen deposition and removal, and a lower maintenance cost of flowers (Ashman and Schoen 1994). Moreover, larger flower longevity can be selected under greater variations of the pollination rate within and between flowering seasons (Xu and Servedio 2021).

However, in many species, flower longevity is plastic, as it is frequently reported that pollen deposition or removal can shorten the flower lifetime, while flower longevity is extended when pollinators are excluded (Primack 1985, Stead 1992, van Doorn and Kamdee 2014, Spigler 2017). The strength of this plasticity of flower longevity also varies across species and populations (Ashman 2004). In many species, pollen deposition at the early stage will greatly decrease flower longevity (Devlin 1984, Yasaka 1998, Arathi 2002, Clayton 1996, Castro 2008, Pacheco 2016). For instance, in *Chlorea alpina*, flowers without pollination open for 8-10 days, while the average longevity of flowers receiving artificial pollen deposition is 3.7 days (Clayton 1996). In contrast, some species show nearly no reaction to pollen deposition, especially in short-lived tropical plants. For example, the average flower longevity of *Lilium philadelphium* is about 9 days with or without pollination exclusion, and no reduced longevity is found after artificial pollen deposition (Edwards 1992). Moreover, hermaphroditic flowers often respond differently to pollen deposition and pollen removal, and the effects of pollen removal on flower longevity is often much weaker than pollen deposition (Clayton 1996, Ishii 2000, Spigler 2017). This pattern remains to be a puzzle, since if hermaphroditic flowers gain fitness through both the male and female functions, early flower senescence should be triggered by either pollinia removal or deposition with a similar degree.

The physiological basis underlying the plasticity of flower longevity is through the regulation of ethylene accumulation (Halevy 1984, Larsen 1995, van Doorn and Kamdee 2014). In many species, flower closure and abscission happen when the ethylene content in the flower tissue reaches a threshold level (Stead 1985). Typically, without pollination, the rate of ethylene production is nearly constant over time, but the rate can be mediated by the amount pollen deposited and removed at a certain period (Stead 1985, Wagstaff 2005). After pollen deposition, the ethylene accumulation rate rises, and this increase has a nearly linear relationship with the amount of pollen deposited (Stead 1985).

Although the physiological mechanism underlying plastic flower longevity is generally clear, the evolutionary causes of the plasticity of flower longevity and how pollination environments affect the strength of flower longevity in the female and male functions remain unclear. The plasticity of flower longevity is expected to evolve under stochastic pollination environments, and can have opposing effects on individual fitness. Specifically, a stronger plasticity of flower longevity induces earlier flower senescence, and thus can be beneficial by saving reproductive resources if the pollination rate fluctuates to be low later. However, if the pollination rate rises to be high later, early senescence can be disadvantageous since although some resources are saved, early senescence may cause a much lower level of pollen deposition and removal compared to longer-lived flowers. Moreover, stochasticity in the pollination environment can happen within or/and between flowering seasons (Arroyo 2013, Duan 2007, Blair 2007, Pacheco 2016), which may have different effects on the evolution of the plasticity of flower longevity (Xu and Servedio 2021).

Another important question about the plasticity of flower longevity is how saved resources increase individual fitness, which remains unclear. Since compared to non-plastic flowers, plastic flowers will have less pollen removed and deposited, the plasticity of flower longevity will not be selected if saved resources through plasticity do not offer any fitness advantage. Here I propose two non-exclusive mechanisms. First, due to increased seed provisioning, saved resources may increase the production of viable offspring through increased fertility or/and viability of offspring. Second, in perennials, saved resources may be stored or allocated to vegetative growth, which increases reproductive resources in the future flowering seasons.

In this study, I investigate how stochasticity within and between flowering seasons can shape the evolution of the plasticity of flower longevity through group or individual selection. To be consistent with previous literature (Ashman and Schoen 1994), I use female and male fitness to refer to the proportion of ovules fertilized and pollen removed in a flower, respectively. The rates of the relative changes in female and male fitness are referred to as female and male fitness accrual rates, respectively. Specifically, I focus on the flowering strategy in 3 dimensions: maximum flower longevity when there is no pollination, strength of plasticity in the female and male functions (i.e., plasticity to ovules fertilization and pollen removal). For group selection, I investigate the optimal strategy that maximizes individual fitness, and for individual selection within a population, I carry out invasion analyses based on the evolutionary game theory (Smith 1982).

Results show that generally, stronger plasticity is selected when female and male fitness accrual rates are higher and more volatile. The evolution of plasticity is also strongly affected by the ratio of the daily maintenance cost of a flower to the construction cost, but its effects on the strength of plasticity depend on the mechanism by which saved resource increase individual fitness. Under group selection, only plasticity in the female function (but not in the male function) tend to be selected, and for the plasticity of flower longevity to be optimal, it requires large fitness advantages through resource saving. Compared to group selection, under individual selection, the evolution of plasticity is more likely, but given the same pollination environment, any combination of the strength of plasticity in female and male functions that adds up to a certain overall strength of plasticity can be evolutionarily stable. Since plastic responses in the male function are rare, the results suggest that the plasticity of flower longevity may evolve mainly through group selection in natural populations.

## Model

I assume a hermaphroditic population, which can be either annual or perennial. For the model presented in this section, I assume the population is outcrossing, but in the result section, the model is extended to account for self-fertilization. The biological meanings of key symbols used in the model are summarized in Table 1. The simulation code is written in R 4.0.2 [available upon acceptance].

**Table 1.**
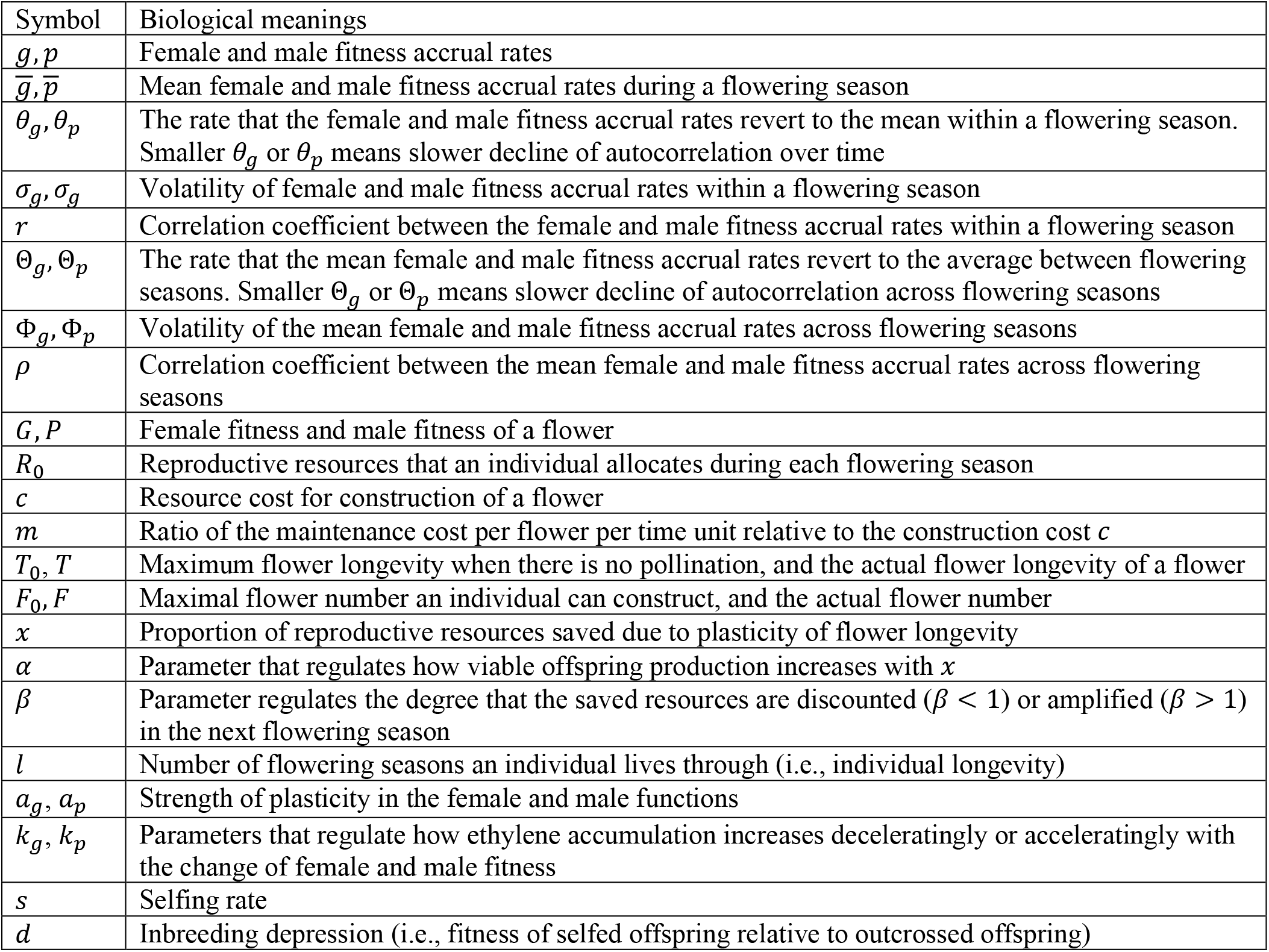
Biological meanings of key symbols used in the model.

| Symbol | Biological meanings |
| --- | --- |
| $g, p$ | Female and male fitness accrual rates |
| $\bar{g}, \bar{p}$ | Mean female and male fitness accrual rates during a flowering season |
| $\theta_g, \theta_p$ | The rate that the female and male fitness accrual rates revert to the mean within a flowering season. Smaller $\theta_g$ or $\theta_p$ means slower decline of autocorrelation over time |
| $\sigma_g, \sigma_p$ | Volatility of female and male fitness accrual rates within a flowering season |
| $r$ | Correlation coefficient between the female and male fitness accrual rates within a flowering season |
| $\Theta_g, \Theta_p$ | The rate that the mean female and male fitness accrual rates revert to the average between flowering seasons. Smaller $\Theta_g$ or $\Theta_p$ means slower decline of autocorrelation across flowering seasons |
| $\Phi_g, \Phi_p$ | Volatility of the mean female and male fitness accrual rates across flowering seasons |
| $\rho$ | Correlation coefficient between the mean female and male fitness accrual rates across flowering seasons |
| $G, P$ | Female fitness and male fitness of a flower |
| $R_0$ | Reproductive resources that an individual allocates during each flowering season |
| $c$ | Resource cost for construction of a flower |
| $m$ | Ratio of the maintenance cost per flower per time unit relative to the construction cost $c$ |
| $T_0, T$ | Maximum flower longevity when there is no pollination, and the actual flower longevity of a flower |
| $F_0, F$ | Maximal flower number an individual can construct, and the actual flower number |
| $x$ | Proportion of reproductive resources saved due to plasticity of flower longevity |
| $\alpha$ | Parameter that regulates how viable offspring production increases with $x$ |
| $\beta$ | Parameter regulates the degree that the saved resources are discounted ($\beta < 1$) or amplified ($\beta > 1$) in the next flowering season |
| $l$ | Number of flowering seasons an individual lives through (i.e., individual longevity) |
| $a_g, a_p$ | Strength of plasticity in the female and male functions |
| $k_g, k_p$ | Parameters that regulate how ethylene accumulation increases deceleratingly or acceleratingly with the change of female and male fitness |
| $s$ | Selfing rate |
| $d$ | Inbreeding depression (i.e., fitness of selfed offspring relative to outcrossed offspring) |

### Pollination process

Following previous models (Ashman and Schoen 1994, Xu and Servedio 2021), for a flower with longevity *T*, the female and male fitness of a flower are

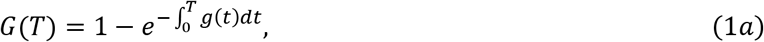

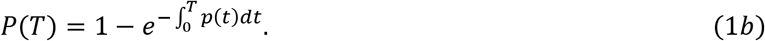

In the above equations, *g*(*t*) and *p*(*t*) are the female and male fitness accrual rates at time *t*. The equations capture the fact that the female or male fitness tend to increase deceleratingly over time and converge to 1.

To model stochastic changes in female and male accrual rates, since pollinator activities are correlated with daily temperature fluctuations, following stochastic models for daily temperature (Benth and Šaltytė-Benth 2005), I assume the changes of female and male fitness accrual rates from time *t* to *t* + *dt* follow two Ornstein–Uhlenbeck processes as

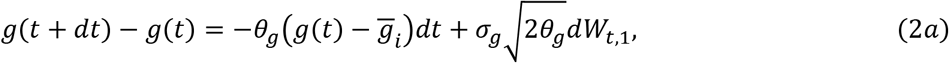

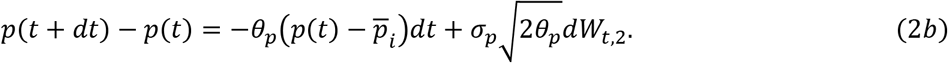

In equations (2a) and (2b), 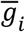 and 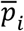 are the expected fitness accrual rates during flowering season *i*. The second term in the right-hand side adds stochasticity, where *W*_*t*,1_ and *W*_*t*,2_ are two Winner processes with a correlation coefficient *r*, since female and male fitness accrual rates may simultaneously fluctuate to be higher or lower. The parameter *σ_g_* characterizes the level of volatility, and the variance of *g*(*t*) is 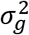. The parameter *θ_g_* is the speed that the female fitness accrual rate reverts to the mean 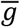, and the autocorrelation between *g*(*t*) and *g*(*t*′) is 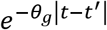. Therefore, a smaller *θ_g_* causes a slower decline of the autocorrelation. Biologically, it means if the rate fluctuates to be low (or high) at some time point, it is more likely to still stay low (or high) later. The above interpretations also apply to the male fitness accrual rate. Since the fitness accrual rates cannot be negative, *g*(*t*) and *p*(*t*) are set to be 0 when their values given by equations (2a) and (2b) are negative. The two stochastic processes are implemented by simulation, and I choose *dt* = 0.1 throughout the results.

For stochasticity between flowering seasons, I assume the mean female and male fitness accrual rates also change as two Ornstein–Uhlenbeck processes as

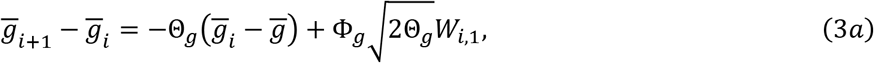

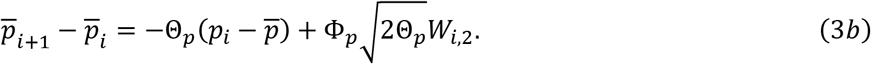

In equations (3a) and (3b), 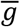 and 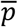 are the expected fitness accrual rates across flowering seasons, and other parameters (Θ*_g_*,Θ*_p_*, Φ_*g*_, Φ*_p_*) have the similar interpretations as those in equations (2a) and (2b). To avoid the situation when 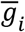 or 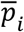 is 0 so that the population becomes extinct, during simulation, 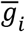 and 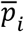 are set to 0.005 when they are smaller than 0.005. When there is stochasticity both within and between flowering seasons, I first simulate the mean fitness accrual rates 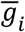 and 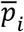 for each flowering season *i*, and within each flowering season, the fitness accrual rates are simulated based on equations (2a) and (2b).

### Flower number, flower longevity, and pollination-induced plasticity

For each flowering season, I assume an individual allocates a fixed amount of reproduction resources *R*_0_, but the actual reproductive resources may be greater than *R*_0_ if there are saved resources from the last flowering season. Due to the resource limitation, there is a trade-off between flower longevity and the flower number. Specifically, I assume each individual produces a constant number of flowers *F*, and flowers open simultaneously with the same longevity. For each flower, the resource cost for construction is *c*, and the maintenance cost per time unit is *mc*, where *m* is the cost of maintenance relative to construction. Therefore, the maximum flower longevity *T*_0_ is

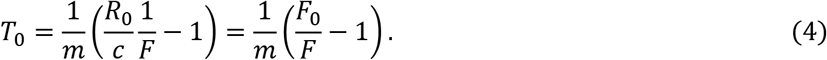

In equation (4), I replace the term *R*_0_/*c* by *F*_0_ to reduce the number of parameters, and *F*_0_ can be interpreted as the maximum number of flowers that an individual can construct. When there is plasticity, the actual flower longevity, denoted by *T*, will be shorter than *T*_0_.

To model pollination-induced plasticity, based on the physiological basis of flower senescence (reviewed in the introduction section), I assume that the relative amount of ethylene accumulated during a small interval from *t* to *t + dt* depends on the changes of the female and male fitness during the interval as

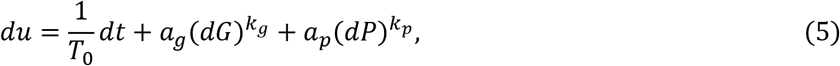

where *dG* and *dP* are the changes of the female and male fitness during the interval, respectively. I assume flower senescence happens when the total amount of ethylene accumulated exceeds a threshold 1. Therefore, after obtaining the female and male fitness accrual rates, the actual flower longevity *T* can be obtained by solving 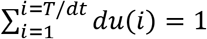, where *du*(*i*) is the amount of ethylene accumulated during the interval from (*i* – 1)*dt* to *idt*. In equation (5), the first term accounts for a constant rate of ethylene production, so that the maximum flower longevity is *T*_0_ when there is no pollination (i.e., *dG* = *dP* = 0). The second and third terms capture increased ethylene production due to changes of the female and male fitness, where *a_g_* and *a_p_* indicate the strength of plasticity in the female and male functions. Since *T*_0_ is a function of the flower number *F* (see equation (4)), the flowering strategy has 3 dimensions {*F, a_g_, a_p_*}. The parameters *k_g_* and *k_p_* regulate how the ethylene accumulation rate increases with changes of the female and male fitness (decelerating when *k_g_, k_p_* < 1; linear when *k_g_ = k_p_* = 1; accelerating when *k_g_, k_p_* > 1). Since *dG, dP* < 1, (*dG*)*^k_g_^* and (*dP*)*^k_p_^* are smaller when *k_g_* and *k_p_* are larger, so *a_g_* and *a_p_* will be larger when *k_g_* and *k_p_* are larger. However, the value of *k_g_* and *k_p_* do not qualitatively change the results because what we are interested in is how factors change the relative values of *a_g_* and *a_p_*. Therefore, throughout the results presented, I set *k_g_ = k_p_* = 0.5. Moreover, it should be noted that equation (5) assumes a linear function, which means that changes of female and male fitness act additively on ethylene accumulation. However, results are qualitatively similar when non-linear functions (e.g., 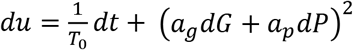) are adopted.

### Fitness advantages through the plasticity of flower longevity

Due to early flower senescence caused by plasticity, the actual resource cost of a flower is *c*(1 + *mT*). Therefore, the proportion of saved resources through plasticity per flower is

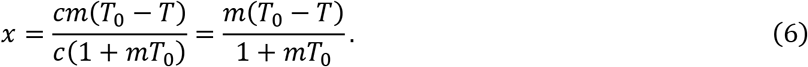

When saved resources increase viable offspring production, I assume the number of viable offspring is proportional to (1 + *x*)^*α*^, where the parameter *α* regulates how viable offspring production increases with *x* (less than a proportional increase when *α* < 1; a proportional increase when *α* = 1; more than a proportional increase when *α* > 1).

When saved resources increase reproductive resources in the next flowering season in perennials, I assume saved resources increase the number of flowers produced and do not affect the maximum flower longevity or the strength of plasticity. Therefore, if the reproductive resources at flowering season *i* is *R_i_*, the flower number at flowering season *i* is 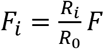. The reproductive resources in flowering season *i* + 1 are

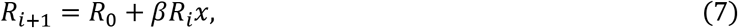

where *R_i_βx* is the total amount of saved resources at flowering season *i*, and *β* is a factor that accounts for either discounting (*β* < 1) or amplification (*β* > 1) of saved resources. Discounting may happen if saved resources are stored (e.g., in the bulb or tuber) and will be partly consumed through respiration, or lost due to inefficiency during energy conversion through synthesis and decomposition. Amplification may happen if saved resources are allocated to vegetative growth, which may produce more resources through photosynthesis.

### Optimal strategy

For group selection, I focus on the optimal flowering strategy that maximizes individual fitness. For annual populations with non-overlapping generations, previous models (Kokko 2020, Xu and Servedio 2021) show that an optimal strategy should maximize the expected fitness in the log scale as

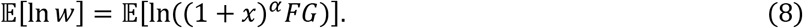

For populations with overlapping generations, I denote the number of flowering seasons an individual lives through by *l*, and assume that the reproductive age is from 1 to *l*. Denote the fitness of an individual during flowering season *i* by *w_i_*, by summing over the offspring produced by individuals at different ages, the total number of offspring produced during flowering season *i, W_i_*, can be obtained by solving the iteration equation

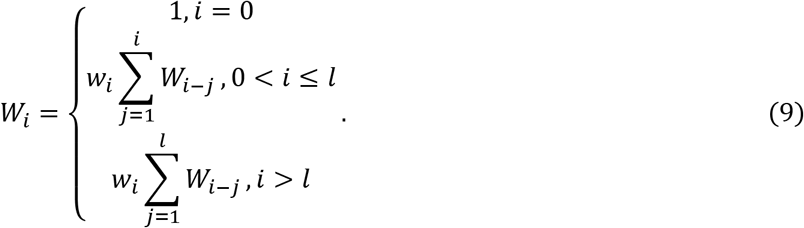

*W_i_* converges to a constant 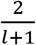 when *w_i_* = 1/*l*, and the population size and the age structure is stabilized. Otherwise, *W_i_* will either converge to 0 when *w_i_* < 1/*l*, or go to infinity when *w_i_* > 1/*l*, so that the population size either declines to 0 or grows to infinite. Therefore, to compare the mean fitness between different flowering strategies, I compare the mean fitness that stabilizes the population *w_e_*, and a larger *w_e_* means higher mean fitness. Since *w_i_* will vary across flowering seasons in the current model due to stochastic pollination environments, to obtain *w_e_*, I replace *w_i_* with 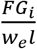, and numerically solve for the value of *w_e_* that makes *W_i_* converges to 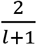 when *i* → ∞.

### Invasion analysis

For the invasion analysis, I consider that a rare mutant individual invades a resident population, and assume flowers of the resident and the mutant open at the same time. I denote the number of flowers, the female and male fitness accrued per flower, and the proportion of saved resources of the resident and the mutant by *F, G, P, x* and *F′, G′, P′, x′*, respectively. However, when the actual flower longevity of the mutant *T′* is larger than that of the resident *T*, the pollen removed after *T* cannot enter the pollen pool to fertilize ovules of the residents. Therefore, the relative contribution of the mutant to the pollen pool is 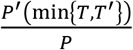, where *P*′(min{*T*, *T*′}) is the male fitness of the mutant at time min{*T*, *T*′}. When saved resources increase viable offspring production, the relative fitness of the mutant is

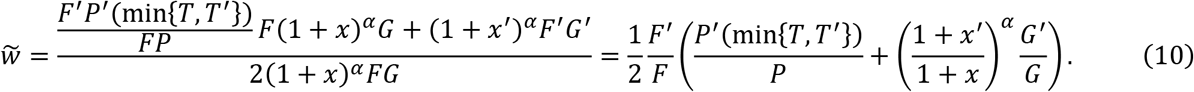

The mutant can invade when 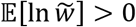. In the middle expression of equation (10), the denominator and the nominator represent the fitness of the resident and the mutant, respectively. The first and second terms in the nominator are the fitness of the mutant obtained through the male and female functions, respectively. The last expression shows that the relative fitness is the arithmetic average of the relative fitness in the male and female functions.

When saved resources increase the reproductive resources in the next flowering season, the invasibility of the mutant can be determined based on equation (9) by replacing the term *w_i_* there with the relative fitness 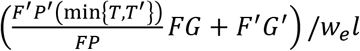, where *w_e_* is the mean fitness of the resident. The mutant population will grow to infinite if its mean relative fitness is higher than 1, and otherwise, the population will decline to 0. Therefore, the mutant can invade if 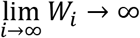 and cannot invade if 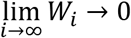.

To obtain the ESS strategy, during the simulation, I assume the mutant strategy differs from the resident strategy at only one dimension with a small step (Δ*F* = ±1, Δ*a_g_* = ±0.01, Δ*a_p_* = ±0.01), since mutations at multiple dimensions are unlikely. The resident strategy is replaced by the mutant strategy once invasion happens. The simulation stops when the population evolves to a strategy that is resistant to the invasion of all 6 possible mutant strategies, and this strategy is considered as an evolutionarily stable strategy (ESS).

## Results

### Optimal strategy

Group selection should select for an optimal flowering strategy that maximizes individual fitness. Since individual fitness only depends on the female fitness but not on the male fitness, we should expect that group selection will not favor plasticity in the male function, and the evolution of plasticity is determined by the female fitness accrual rate. Therefore, we can always set *a_p_* = 0 in equation (5), and the strategy space is reduced to 2 dimensions (i.e., {*F, a_g_*}). In fact, even in the 3-dimensional space {*F, a_g_, a_p_*}, simulations show that on optimality, *a_p_* = 0. The only exception is when the female and male fitness accrual rates are highly correlated, but in this case, only an overall strength of plasticity *α* = *a_g_* + *a_p_* can be determined, and any combination of *a_g_* and *a_p_* satisfying *a_g_* + *a_p_* = *α* is an optimal strategy.

Fig. 1 shows how individual fitness changes with the flower number or the strength of plasticity. As Fig. 1(a) shows, given a fixed strength of plasticity, there is a certain flower number (thus certain maximum flower longevity *T*_0_) that maximizes individual fitness, and this optimal flower number is smaller when the plasticity is stronger, since a stronger plasticity should favor larger maximum flower longevity. Similarly, Fig. 1(b) shows that given a fixed flower number, a certain strength of plasticity confers the highest individual fitness, and this optimal level of plasticity is lower when the flower number is larger (thus shorter *T*_0_).

**Figure 1.**
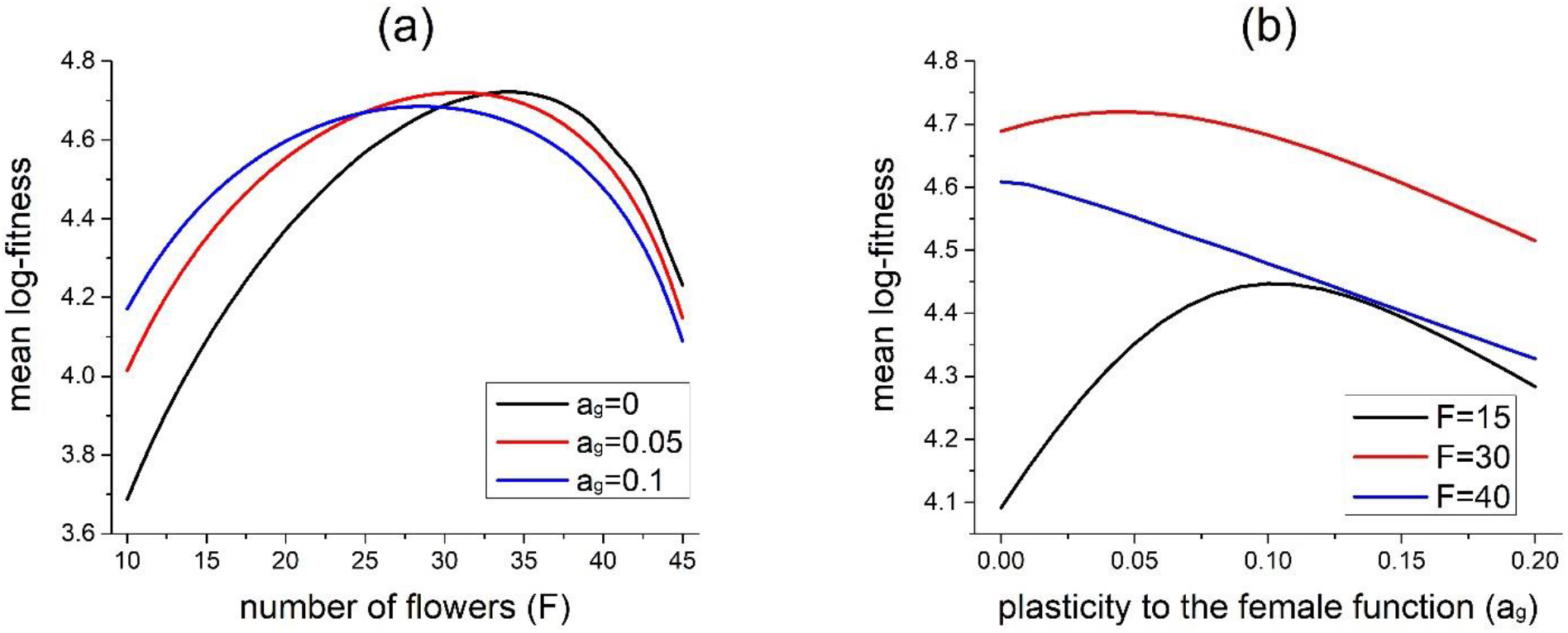
Effects of the flower number (*F*) and the level of plasticity in the female function (*a_g_*) on the mean log-fitness. Parameters are *F*_0_ = 50, *m* = 0.1, *l* = 20, *α* = 20, 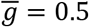, *θ_g_* = 0.1, 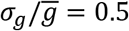. Similar results are found when the saved resources are sued to increase the reproductive resources in the next year in perennials.

By allowing both the flower number and the strength of plasticity to change, Tables 2 and 3 show how different factors affect the optimal strategy. However, the plasticity of flower longevity is likely to be optimal only when the fitness advantage through resource saving is large. Specifically, when saved resources increase viable offspring production, the strength of plasticity on optimality is not 0 only when *α* > 1, which means viable offspring production increases acceleratingly with the saved resources. When saved resources increase reproductive resources in the next flowering season, plasticity may be optimal only when *β* > 1, which means reproductive resources are amplified and thus accumulate over flowering seasons as a snow-ball effect. Moreover, under this case, the fitness advantage of plasticity through resource saving is greater when individuals can live longer (larger *l*). Therefore, the plasticity on optimality is stronger when *l* is larger, as shown in Table 3.

**Table 2.** The optimal strategy {*F, a_g_*} (*a_p_* = 0 on optimality) when saved resources increase viable offspring production under stochasticity within flowering seasons. Unless specified, *m* = 0.05, *F*_0_ = 50, *α* = 2, *k_g_* = 0.5.

| | $\bar{g} = 0.2$ | $\bar{g} = 0.5$ | $\bar{g} = 0.8$ | $\alpha = 3, \bar{g} = 0.2$ | $m = 0.2$ | $F_0 = 100$ |
| --- | --- | --- | --- | --- | --- | --- |
| $\theta_g = 1, \sigma_g/\bar{g} = 0.2$ | 34, 0.00 | 39, 0.01 | 42, 0.00 | 25, 0.06 | 30, 0.00 | 79, 0.00 |
| $\theta_g = 0.01, \sigma_g/\bar{g} = 0.2$ | 34, 0.00 | 39, 0.01 | 42, 0.00 | 25, 0.06 | 30, 0.00 | 79, 0.00 |
| $\theta_g = 1, \sigma_g/\bar{g} = 0.5$ | 34, 0.00 | 39, 0.01 | 41, 0.01 | 25, 0.06 | 29, 0.01 | 77, 0.01 |
| $\theta_g = 0.01, \sigma_g/\bar{g} = 0.5$ | 31, 0.02 | 36, 0.03 | 39, 0.04 | 24, 0.06 | 27, 0.03 | 73, 0.03 |
| $\theta_g = 1, \sigma_g/\bar{g} = 0.8$ | 34, 0.00 | 39, 0.01 | 41, 0.02 | 26, 0.06 | 29, 0.01 | 78, 0.01 |
| $\theta_g = 0.01, \sigma_g/\bar{g} = 0.8$ | 29, 0.02 | 34, 0.04 | 36, 0.04 | 23, 0.06 | 25, 0.04 | 67, 0.04 |

**Table 3.** The optimal strategy {*F, a_g_*} when saved resources increase the reproductive resources in the next flowering season under stochasticity within flowering seasons. Unless specified, *m* = 0.05, *F*_0_ = 50, *l* = 10, *β* = 1.2, *k_g_* = 0.5, 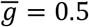.

| | $\bar{g} = 0.2$ | $\bar{g} = 0.5$ | $\bar{g} = 0.8$ | $l = 5$ | $l = 20$ | $m = 0.2$ | $F_0 = 100$ |
| --- | --- | --- | --- | --- | --- | --- | --- |
| $\theta_g = 1, \sigma_g/\bar{g} = 0.2$ | 12, 0.10 | 15, 0.14 | 16, 0.17 | 39, 0.00 | 10, 0.16 | 11, 0.16 | 30, 0.14 |
| $\theta_g = 0.01, \sigma_g/\bar{g} = 0.2$ | 13, 0.10 | 15, 0.14 | 16, 0.17 | 39, 0.00 | 10, 0.16 | 12, 0.16 | 30, 0.14 |
| $\theta_g = 1, \sigma_g/\bar{g} = 0.5$ | 13, 0.10 | 15, 0.14 | 16, 0.17 | 39, 0.00 | 10, 0.16 | 11, 0.17 | 30, 0.13 |
| $\theta_g = 0.01, \sigma_g/\bar{g} = 0.5$ | 13, 0.10 | 15, 0.14 | 16, 0.18 | 39, 0.00 | 10, 0.17 | 11, 0.17 | 30, 0.14 |
| $\theta_g = 1, \sigma_g/\bar{g} = 0.8$ | 13, 0.11 | 15, 0.15 | 16, 0.19 | 39, 0.01 | 10, 0.17 | 12, 0.18 | 30, 0.14 |
| $\theta_g = 0.01, \sigma_g/\bar{g} = 0.8$ | 12, 0.11 | 15, 0.15 | 16, 0.19 | 37, 0.02 | 10, 0.18 | 11, 0.17 | 30, 0.15 |

Generally, a stronger plasticity is likely to be optimal when the mean female fitness accrual rate 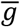 and volatility *σ_g_* are higher. Nevertheless, a higher mean fitness accrual rate not only favors a stronger plasticity, but also favors smaller maximum flower longevity *T*_0_ (compare results at 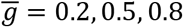 in Tables 2 and 3). This is an important point, since it means if variation in the strength of plasticity is mainly due to variation in the mean fitness accrual rate, shorter-lived flowers will exhibit stronger plastic response. However, given the same mean fitness accrual rates, stronger plasticity is associated with a smaller flower numbers and thus larger maximum flower longevity. The optimal strength of plasticity is also affected by the ratio of the maintenance cost of a flower relative to the construction cost, *m*, but its effects depend on how the saved resources increase individual fitness. Specifically, on optimality, a larger *m* reduces the strength of plasticity when saved resources increase viable offspring production (compare results in the column 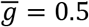 and *m* = 0.2 in Table 2), but increases the strength of optimal plasticity when saved resources increase reproductive resources in the next flowering season (compare results in the column 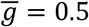 and *m* = 0.2 in Table 3).

### Invasion analysis

Under individual selection, the evolution of the plasticity of flower longevity is more likely than that under group selection. Specifically, under individual selection, plasticity can be an ESS even when *α* < 1 or *β* < 1, and given the same fitness advantage through resource saving, the plasticity at the ESS is much stronger than the plasticity at the optimal strategy under group selection (e.g., compare Table 2 and Table 4). This is partly because the invasion of a mutant with shorter maximum flower longevity or stronger plasticity is more likely than the invasion of a mutant with larger maximum flower longevity or weaker plasticity. Specifically, a mutant with larger maximum flower longevity or weaker plasticity tends to have longer-lived flowers than the resident, and thus not all its removed pollen can enter the pollen pool to fertilize ovules of the residents. This factor is captured by the term 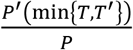 in equation (9). As a consequence, the evolution of maximum flower longevity and the strength of plasticity will depend on the initial strategy of the population and the invasion is generally unidirectional. Specifically, a population starting with individuals with few long-lived flowers and weak plasticity is susceptible to the invasion of mutants with stronger plasticity or smaller maximum flower longevity. In contrast, if initially, individuals have many short-lived flowers and strong plasticity, even the strength of plasticity is excessively high, which does not maximize female and male fitness, the population can still resist the invasion of mutants with weaker plasticity. Therefore, in Table 4, results presented are the ESS that a population starting with long-lived, non-plastic flowers evolves towards. However, given the same parameters, a strategy with a stronger plasticity or larger flower numbers than the result shown in Table 4 may also be an ESS.

**Table 4.**
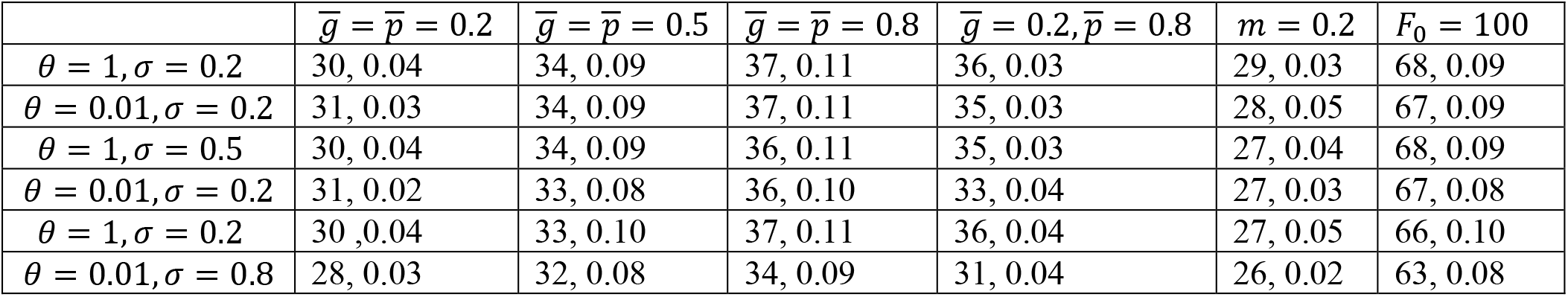
The ESS {*F, a_g_* + *a_p_*} when saved resources increase viable offspring production under stochasticity within flowering seasons. Unless specified, *m* = 0.05, *F*_0_ = 50, *α* = 2, *k_g_* = *k_p_* = 0.5, 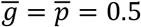, and *θ* = *θ_g_* = *θ_p_*, 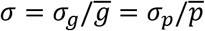. The initial strategy of the population is *F* = 5, *a_g_* = *a_p_* = 0.

Another important point is that under individual selection, stochastic pollination environments can only select for a certain strength of the overall plasticity *a = a_g_ + a_f_*. The specific combination of the strength of plasticity in the female and male functions (*a_g_, a_p_*) cannot be determined because any combination satisfying *a_g_ + a_f_ = a* will be evolutionarily stable. This is true even when the mean fitness accrual rates or the strength of volatility greatly differ between the female and male functions. In other words, when the female fitness accrual rate is high and volatile, while the male fitness accrual rate is low and more constant, a strategy with strong plasticity in the male function and weak plasticity in the female function can still be an ESS. Mathematically, this result means that the ESS is not a point in the 3-dimensional strategy space, but lies along a 1-dimensional line. Therefore, the strength of plasticity at the ESS in the female and male functions depends on the initial condition and the invasion history. Although it is suspected that this result may be due to the assumption of additivity in equation (5) when modelling plasticity, similar results are found even when non-linear functions are adopted.

The above results hold for both stochasticity within and between flowering seasons, and do not depend on whether saved resources increase viable offspring production or reproductive resources in the next flowering season. However, the time scale of stochasticity and the mechanism by which saved resources increase individual fitness do affect the effects of some parameters (e.g., autocorrelation, maximum flower number) on the strength of the ESS plasticity, as summarized in Table 5. Generally, the strength of the ESS plasticity is mainly determined by the mean female and male fitness accrual rates 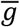 and 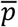, as well as the maintenance cost of flowers relative to construction, *m*. Similar to the results under group selection, a smaller flower number (thus shorter maximum flower longevity) and stronger plasticity are selected when 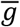 and 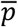 are higher (see Table 4). Interestingly, when mean fitness accrual rates differ in female and male functions, the ESS plasticity is constraint by the function with a lower mean fitness accrual rate. For example, in Table 4, when 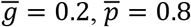, the strength of the ESS plasticity is similar to that when 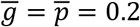, and is much smaller than that when 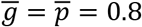. However, this effect is absent when saved resources increase reproductive resources in the next flowering season and there is stochasticity between flowering seasons (see Table 5).

**Table 5.** A summary of the effects of factors on the evolution of plasticity based on Tables 1–4 and S1-S10 (the biological meanings of the parameters can be found in Table 1). The symbol “+” and “–” indicate whether a larger value of the parameter promotes or inhibits the evolution of plasticity, and “+/–” means both cases may happen. The word “constraint” means the strength of plasticity at the ESS is determined by the function with a lower mean fitness accrual rate. Under group selection, only plasticity in the female function *a_g_* will be favored on optimality. Under individual selection, only an overall strength of plasticity can be determined. Generally, the autocorrelation and volatility of the pollination environment have only small effects on the optimal or the evolutionarily stable strategy. Also, the maximum number of flowers *F*_0_ and the correlation coefficient between female and male fitness accrual rates almost have no effect (see Table S10).

| Level of selection | Fitness advantage of resource saving | Stochasticity within flowering seasons | Stochasticity between flowering seasons | Stochasticity at both time scales |
| --- | --- | --- | --- | --- |
| Group selection (optimal strategy) | Increased viable offspring production | $\bar{g}$: +<br>$\sigma_g$: +<br>$\theta_g$: -<br>$m$: - | $\bar{g}$: +<br>$\Phi_g$: +<br>$\Theta_g$: +<br>$m$: - | N/A |
| | Increased reproductive resources in the next flowering season | $\bar{g}$: +<br>$\sigma_g$: +<br>$\theta_g$: +<br>$m$: +<br>$l$: + | $\bar{g}$: +<br>$\Phi_g$: +<br>$\Theta_g$: +/-<br>$m$: +<br>$l$: + | N/A |
| Individual selection (evolutionarily stable strategy) | Increased viable offspring production | $\bar{g}, \bar{p}$: +<br>$\sigma_g, \sigma_p$: +<br>$\theta_g, \theta_p$: +<br>$m$: -<br>constraint | $\bar{g}, \bar{p}$: +<br>$\Phi_g, \Phi_p$: +<br>$\Theta_g, \Theta_p$: +/-<br>$m$: -<br>constraint | constraint |
| | Increased reproductive resources in the next flowering season | $\bar{g}, \bar{p}$: +<br>$\sigma_g, \sigma_p$: +<br>$\theta_g, \theta_p$: +<br>$m$: +<br>$l$: +<br>constraint | $\bar{g}, \bar{p}$: +<br>$\Phi_g, \Phi_p$: +<br>$\Theta_g, \Theta_p$: +<br>$m$: +<br>$l$: +<br>not constraint | not constraint |

### Effects of selfing

The previous results assume the population is outcrossing, but selfing is quite common in hermaphroditic plants. The previous study suggests that different modes of selfing may have different effects on the evolution of flower longevity (Xu 2021). Therefore, here I consider two cases of selfing, depending on whether selfing relies on the pollination process. When selfing does not depend on the pollination process (e.g., prior selfing, production of a combination of chasmogamous and cleistogamous flowers), I denote the proportion of selfed ovules out of the total ovules (either fertilized or unfertilized) by s. Therefore, the female fitness of a flower with flower longevity *T* is

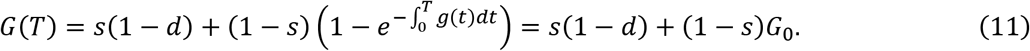

In equation (11), *d* is inbreeding depression, measured as the fitness of selfed offspring relative to outcrossed offspring (Charlesworth and Willis 2009), and *G*_0_ is the female fitness when the flower is completely outcrossing. The first and second terms in the middle expression are the fitness obtained through selfing and outcrossing, respectively, after accounting for inbreeding depression. When selfing depends on the pollination process (e.g., competing selfing, geitonogamy), I assume a proportion s of fertilized ovules is self-fertilized, and the strength of plasticity does not depend on the source of pollen (as found in Spigler 2017). Therefore, the female fitness of a flower is

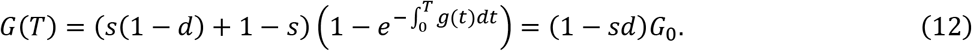

It can be seen that selfing discounts the change of the female fitness by a factor (1 – *s*) in equation (11), and (1 – *sd*) in equation (12). Based on equation (5), given the same pollination history, an outcrossing flower with strategy {*F, a_g_, a_p_*} will have the same actual flower longevity as a selfing flower with strategy 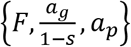 (for equation (11)) or 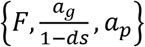 (for equation (12)). In the Appendix, it is shown that under group selection, selfing tends to increase the strength of plasticity on optimality. For individual selection within a population, when selfing does not depend on the pollination process, selfing should inhibit the invasion of plasticity and thus reduce the strength of the ESS plasticity, since selfing reduces the fitness advantage through the female function. However, when selfing depends on the pollination process, it should have no effect on the ESS plasticity.

## Discussion

This study investigates the evolution of the maximum flower longevity and the plasticity of flower longevity under stochastic pollination environments through group and individual selection. The model considers two possible mechanisms by which saved resource through the plasticity of flower longevity increase individual fitness: increased viable offspring production, or increased reproductive resources in the next flowering season in perennials. The key results are summarized in Table 5. Generally, a stronger plasticity tends to be optimal or evolutionarily stable when mean female and male fitness accrual rates are higher and the rates are more volatile. The strength of plasticity is also greatly affected by the daily maintenance cost of flowers relative to construction, but like other factors (e.g., autocorrelation), its effects depend on the time scale of stochasticity and the mechanism by which saved resources increase individual fitness (see Table 5).

Contrary to the common expectation that the plasticity of flower longevity should be more likely to evolve in longer-lived flowers, results show that higher mean female and male fitness accrual rates not only select for stronger plasticity, but also shorter maximum flower longevity. Nevertheless, given fixed mean fitness accrual rates, stronger plasticity is associated with a smaller flower number and thus larger maximum flower longevity, which is consistent with the expectation. Since pollination-induced senescence is concentrated in families with long-lived flowers, and absent in groups with very short-lived flowers (Stead 1992, Van Doom 1997), these results suggest that differences in the strength of plasticity of flower longevity across species may be not mainly due to differences in the mean fitness accrual rates, but due to differences in other factors, such as autocorrelation and volatility in the pollination environment, the maintenance cost of flowers relative to construction, and individual longevity. Specifically, the result that stronger plasticity tends to evolve under stronger volatility may explain why tropical species exhibit less plasticity of flower longevity (van Doom 1997), as well as an increased plasticity of flower longevity in alpine plant populations from high elevation compared to low elevation populations (Trunschke and Stöcklin 2017). The result may also explain why in many orchids, flowers longevity is plastic, since pollen aggregation increases volatility of the male fitness accrual rate.

This study also shows that the evolution of plasticity of flower longevity is easier through individual selection than group selection. Under group selection, for plasticity to be the optimal strategy, it requires that saved resources either cause a more than proportional increase of viable offspring production, or are amplified in the next flowering season. In contrast, under individual selection, stronger plasticity can evolve even under small fitness advantages through resource saving. Also, within a population, the evolution of plasticity tends to be unidirectional. This is because compared to a mutant with stronger plasticity, a mutant with weaker plasticity is less likely to invade since it will have longer-lived flowers than the resident, so that not all its removed pollen can fertilize the ovules of the residents. Therefore, there will be multiple evolutionarily stable strategies (ESS), and the ESS that a population will evolve to depend on the initial condition and the invasion history.

Despite the theoretical prediction that the plasticity of flower longevity is more likely to evolve through individual selection, the fact that plastic responses to the change of the male function (i.e., pollen removal) in plants with hermaphroditic flowers are rare suggest that the plasticity of flower longevity may be mainly selected through group selection. This is because under group selection, only plasticity in the female function will be favored, while plasticity in the male function will not be optimal in maximizing individual fitness, since individual fitness does not depend on the male fitness under group selection. Although plasticity in the male function can be selected under individual selection, given the same pollination environment, any combination of the strength of plasticity in the female and male functions that adds up to a certain overall strength of plasticity can be evolutionarily stable. In other words, even under a pollination environment with very volatile ovule fertilization rate but more constant pollen removal rate, a flowering strategy with a strong plasticity to pollen removal and no plasticity to ovule fertilization can be evolutionarily stable.

Despite the numerous evidence on pollination-induced plasticity, detailed knowledges about the plasticity of flower longevity are still lacking. Most studies deposit a large amount of pollen at the early stage of anthesis to show the existence of plasticity, but only few studies investigate how flower longevity changes with the amount of pollen deposited and removed, as well as the source of pollen (e.g., Spigler 2017). Also, the model assumes that the strength of plasticity is constant, but the strength of plasticity of a flower may change over time. Therefore, future studies that apply pollen deposition with different amounts at different time points during the lifetime of a flower may offer more insights for our understanding of plasticity of flower longevity. Moreover, the results also suggest it may be important to investigate how and to what extent can saved resources through the plasticity of flower longevity increase individual fitness.

Finally, it should be emphasized that the current study focuses on pollination-induced plasticity of flower longevity, but plastic responses of flower longevity to abiotic factors (e.g., temperature) are also reported (Yasaka et al. 1998, Arroyo et al. 2013). In fact, meta-analysis shows that the strength of selection by abiotic factors may be comparable to pollination-mediated selection (Caruso et al. 2019). Therefore, the evolution of the plasticity of flower longevity may be also mediated by temporal fluctuations of abiotic factors, which tend to affect the relative maintenance cost of flowers and the amount of reproductive resources. In short, this study examines that how the plasticity of flower longevity in the female and male functions is shaped by stochasticity in the pollination environment at different time scales and selection at different levels.

## Acknowledgments

I would like to thank my parents for shaping my interest in flowering strategies. This work was supported by the National Science Foundation (Grant No. DEB 1939290).

## Appendix

Here I qualitatively analyze the effects of selfing on the evolution of the plasticity of flower longevity under group and individual selection.

### Optimal strategy under group selection

Under group selection, on optimality, the strength of plasticity in the male function is *a_p_* = 0, so the strategy space is reduced to 2 dimensions {*F, a_g_*}. The log-fitness of an outcrossing and selfing individual are

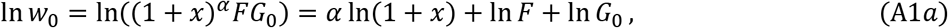

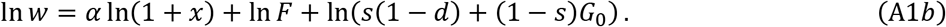

When selfing does not depend on the pollination process, given the same pollination history, equation (11) shows that an outcrossing flower with strategy {*F, a_g_*} will have the same actual flower longevity as a selfing flower with strategy 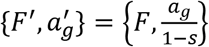, and thus the same proportion of resources saved *x*. If strategy {*F, a_g_*} maximizes the mean fitness of an outcrossing individual, 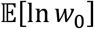, the first-order conditions require

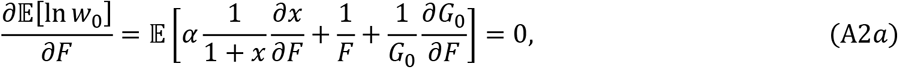

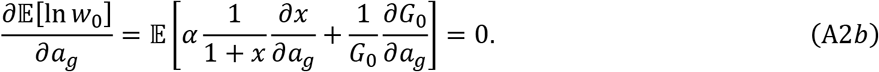

The first-order conditions for a selfing individual with strategy 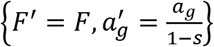 are

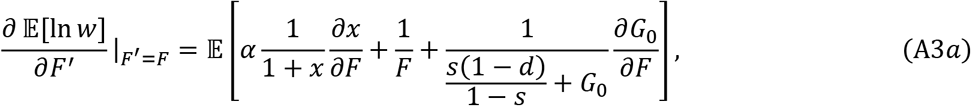

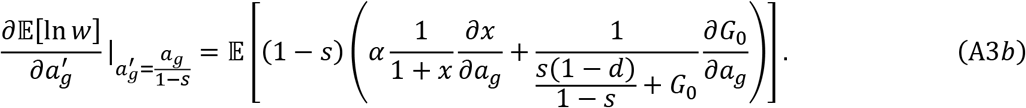

Since a larger flower number *F* or a stronger plasticity *a_g_* reduces the actual flower longevity (thus lowers female fitness), 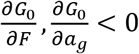. Therefore, 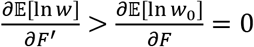, and 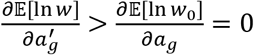. This means that for a selfing individual, on optimality, *F*′ > *F* and 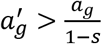.

When selfing depends on the pollination process, given the same pollination history, based on equation (12), the fitness of a selfing individual is

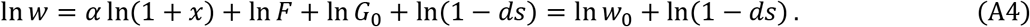

If strategy {*F*, *a_g_*} maximizes the expected log-fitness of an outcrossing individual 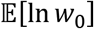, since under selfing, strategy 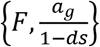 gives the same actual flower longevity (and thus the same *G*_0_), it also maximizes the expected log-fitness of a selfing individual 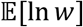. Therefore, selfing increases the strength of plasticity.

### Relative fitness under individual selection

Here I assume the selfing rates of the mutant and the resident are the same. Under individual selection, selfing not only affects the female fitness, but may also affect the fitness obtained through the male function because selfing reduces the number of ovules that the mutant can fertilize through pollen dispersal. When selfing does not depend on the pollination process, the relative fitness given in equation (10) should be modified to

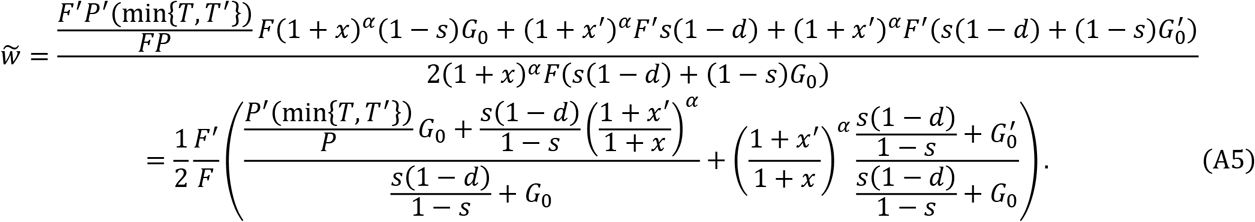

It is clear that 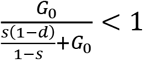, and it can be proved that 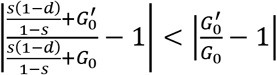. Therefore, selfing reduces the relative fitness through both the male and female functions, which should inhibit the invasion of plasticity. When selfing depends on the pollination process, based on equation (12), the relative fitness is

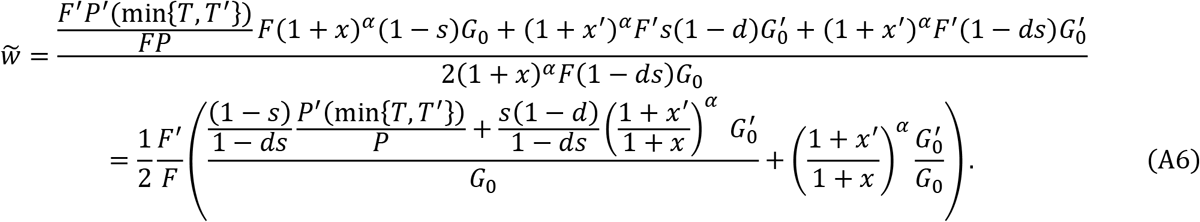

Therefore, selfing will increase the relative fitness through the male function but does not affect the relative fitness through the female function.

## Supplementary Tables and Figures

**Table S1.** The optimal strategy when saved resources increase the viable offspring production under stochasticity between flowering seasons. Unless specified, *m* = 0.05, *F*_0_ = 50, *α* = 2, *k_g_* = 0.5, 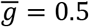.

| | $\bar{g} = 0.2$ | $\bar{g} = 0.5$ | $\bar{g} = 0.8$ | $m = 0.2$ | $F_0 = 100$ |
| --- | --- | --- | --- | --- | --- |
| $\Theta_g = 1, \Phi_g/\bar{g} = 0.2$ | 34, 0.00 | 39, 0.01 | 42, 0.00 | 35, 0.00 | 79, 0.00 |
| $\Theta_g = 0.01, \Phi_g/\bar{g} = 0.2$ | 33, 0.00 | 39, 0.01 | 42, 0.00 | 35, 0.01 | 78, 0.00 |
| $\Theta_g = 1, \Phi_g/\bar{g} = 0.5$ | 29, 0.03 | 34, 0.05 | 36, 0.07 | 30, 0.04 | 68, 0.05 |
| $\Theta_g = 0.01, \Phi_g/\bar{g} = 0.5$ | 34, 0.01 | 36, 0.03 | 41, 0.02 | 32, 0.02 | 76, 0.02 |
| $\Theta_g = 1, \Phi_g/\bar{g} = 0.8$ | 28, 0.04 | 31, 0.07 | 32, 0.10 | 28, 0.07 | 62, 0.07 |
| $\Theta_g = 0.01, \Phi_g/\bar{g} = 0.8$ | 30, 0.02 | 37, 0.03 | 39, 0.05 | 31, 0.03 | 74, 0.03 |

**Table S2.**
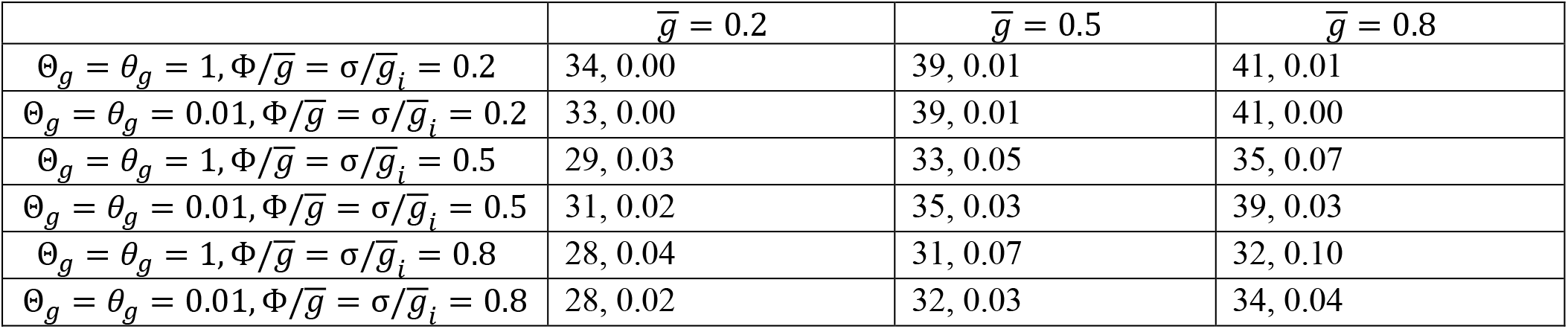
The optimal strategy when saved resources increase the viable offspring production under stochasticity both within and between flowering seasons. Unless specified, *m* = 0.05, *F*_0_ = 50, *α* = 2, *k_g_* = 0.5.

| | $\bar{g} = 0.2$ | $\bar{g} = 0.5$ | $\bar{g} = 0.8$ |
| --- | --- | --- | --- |
| $\Theta_g = \theta_g = 1, \Phi/\bar{g} = \sigma/\bar{g}_i = 0.2$ | 34, 0.00 | 39, 0.01 | 41, 0.01 |
| $\Theta_g = \theta_g = 0.01, \Phi/\bar{g} = \sigma/\bar{g}_i = 0.2$ | 33, 0.00 | 39, 0.01 | 41, 0.00 |
| $\Theta_g = \theta_g = 1, \Phi/\bar{g} = \sigma/\bar{g}_i = 0.5$ | 29, 0.03 | 33, 0.05 | 35, 0.07 |
| $\Theta_g = \theta_g = 0.01, \Phi/\bar{g} = \sigma/\bar{g}_i = 0.5$ | 31, 0.02 | 35, 0.03 | 39, 0.03 |
| $\Theta_g = \theta_g = 1, \Phi/\bar{g} = \sigma/\bar{g}_i = 0.8$ | 28, 0.04 | 31, 0.07 | 32, 0.10 |
| $\Theta_g = \theta_g = 0.01, \Phi/\bar{g} = \sigma/\bar{g}_i = 0.8$ | 28, 0.02 | 32, 0.03 | 34, 0.04 |

**Table S3.** The optimal strategy when saved resources increase the reproductive resources in the next flowering season under stochasticity between flowering seasons. Unless specified, *m* = 0.05, *F*_0_ = 50, *l* = 10, *β* = 1.2, *k_g_* = 0.5, 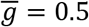.

| | $\bar{g} = 0.2$ | $\bar{g} = 0.5$ | $\bar{g} = 0.8$ | $l = 5$ | $m = 0.2$ | $F_0 = 100$ |
| --- | --- | --- | --- | --- | --- | --- |
| $\Theta_g = 1, \Phi_g/\bar{g} = 0.2$ | 14, 0.14 | 15, 0.15 | 16, 0.17 | 39, 0.01 | 11, 0.17 | 30, 0.14 |
| $\Theta_g = 0.01, \Phi_g/\bar{g} = 0.2$ | 14, 0.13 | 15, 0.15 | 16, 0.17 | 39, 0.01 | 11, 0.16 | 29, 0.14 |
| $\Theta_g = 1, \Phi_g/\bar{g} = 0.5$ | 14, 0.13 | 15, 0.15 | 16, 0.17 | 39, 0.01 | 18, 0.14 | 31, 0.15 |
| $\Theta_g = 0.01, \Phi_g/\bar{g} = 0.5$ | 14, 0.13 | 15, 0.15 | 16, 0.17 | 39, 0.01 | 11, 0.16 | 31, 0.15 |
| $\Theta_g = 1, \Phi_g/\bar{g} = 0.8$ | 14, 0.13 | 15, 0.15 | 16, 0.17 | 39, 0.01 | 33, 0.00 | 50, 0.14 |
| $\Theta_g = 0.01, \Phi_g/\bar{g} = 0.8$ | 14, 0.13 | 15, 0.15 | 16, 0.17 | 39, 0.01 | 12, 0.19 | 31, 0.16 |

**Table S4.** The optimal strategy when saved resources increase the reproductive resources in the next flowering season under stochasticity both within and between flowering seasons. Unless specified, *m* = 0.05, *F*_0_ = 50, *β* = 1.2, *k_g_* = 0.5.

| | $\bar{g} = 0.2$ | $\bar{g} = 0.5$ | $\bar{g} = 0.8$ |
| --- | --- | --- | --- |
| $\Theta_g = \theta_g = 1, \Phi/\bar{g} = \sigma/\bar{g}_i = 0.2$ | 13, 0.10 | 15, 0.14 | 16, 0.17 |
| $\Theta_g = \theta_g = 0.01, \Phi/\bar{g} = \sigma/\bar{g}_i = 0.2$ | 13, 0.10 | 15, 0.14 | 16, 0.17 |
| $\Theta_g = \theta_g = 1, \Phi/\bar{g} = \sigma/\bar{g}_i = 0.5$ | 13, 0.11 | 16, 0.16 | 17, 0.19 |
| $\Theta_g = \theta_g = 0.01, \Phi/\bar{g} = \sigma/\bar{g}_i = 0.5$ | 13, 0.11 | 16, 0.15 | 17, 0.19 |
| $\Theta_g = \theta_g = 1, \Phi/\bar{g} = \sigma/\bar{g}_i = 0.8$ | 35, 0.00 | 40, 0.00 | 41, 0.00 |
| $\Theta_g = \theta_g = 0.01, \Phi/\bar{g} = \sigma/\bar{g}_i = 0.8$ | 13, 0.11 | 17, 0.16 | 20, 0.18 |

**Table S5.** The ESS {*F, a_g_ + a_p_*} when saved resources increase viable offspring production under stochasticity between flowering seasons. Unless specified, *m* = 0.05, *F*_0_ = 50, *α* = 2, *k_g_* = *k_p_* = 0.5, 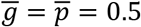, and Θ = Θ*_g_* = Θ*_p_*, 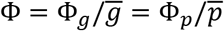. The initial strategy is *F* = 5, *a_g_* = *a_p_* = 0.

| | $\bar{g} = \bar{p} = 0.2$ | $\bar{g} = \bar{p} = 0.5$ | $\bar{g} = \bar{p} = 0.8$ | $\bar{g} = 0.2, \bar{p} = 0.8$ | $m = 0.2$ | $F_0 = 100$ |
| --- | --- | --- | --- | --- | --- | --- |
| $\Theta = 1, \Phi = 0.2$ | 30, 0.04 | 35, 0.07 | 37, 0.11 | 30, 0.04 | 28, 0.04 | 68, 0.09 |
| $\Theta = 0.01, \Phi = 0.2$ | 30, 0.04 | 34, 0.08 | 37, 0.10 | 30, 0.04 | 29, 0.02 | 68, 0.09 |
| $\Theta = 1, \Phi = 0.5$ | 28, 0.04 | 34, 0.06 | 36, 0.10 | 29, 0.04 | 27, 0.05 | 63, 0.09 |
| $\Theta = 0.01, \Phi = 0.5$ | 29, 0.04 | 35, 0.08 | 36, 0.12 | 29, 0.04 | 27, 0.04 | 68, 0.10 |
| $\Theta = 1, \Phi = 0.8$ | 29, 0.03 | 31, 0.07 | 37, 0.11 | 28, 0.04 | 26, 0.06 | 60, 0.09 |
| $\Theta = 0.01, \Phi = 0.8$ | 29, 0.05 | 33, 0.08 | 34, 0.10 | 28, 0.04 | 28, 0.06 | 66, 0.09 |

**Table S6.** The ESS {*F, a_g_ + a_p_*} when saved resources increase viable offspring production under stochasticity both within and between flowering seasons. Unless specified, *m* = 0.05, *F*_0_ = 50, *α* = 2, *k_g_* = *k_p_* = 0.5, 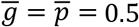. Also, *θ* = *θ_g_* = *θ_p_*, 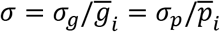, Θ*_g_* = Θ*_p_* = Θ and 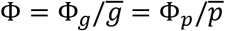. The initial strategy is *F* = 5, *a_g_* = *a_p_* = 0.

| | $\bar{g} = \bar{p} = 0.2$ | $\bar{g} = \bar{p} = 0.5$ | $\bar{g} = \bar{p} = 0.8$ | $\bar{g} = 0.2, \bar{p} = 0.8$ |
| --- | --- | --- | --- | --- |
| $\Theta = \theta = 1, \Phi = \sigma = 0.2$ | 29, 0.05 | 34, 0.08 | 36, 0.11 | 35, 0.03 |
| $\Theta = \theta = 0.01, \Phi = \sigma = 0.2$ | 29, 0.05 | 34, 0.09 | 37, 0.11 | 36, 0.02 |
| $\Theta = \theta = 1, \Phi = \sigma = 0.5$ | 29, 0.04 | 32, 0.08 | 34, 0.11 | 32, 0.04 |
| $\Theta = \theta = 0.01, \Phi = \sigma = 0.5$ | 28, 0.04 | 33, 0.07 | 34, 0.10 | 32, 0.04 |
| $\Theta = \theta = 1, \Phi = \sigma = 0.8$ | 28, 0.04 | 31, 0.09 | 32, 0.10 | 30, 0.06 |
| $\Theta = \theta = 0.01, \Phi = \sigma = 0.8$ | 27, 0.04 | 30, 0.06 | 33, 0.08 | 30, 0.05 |

**Table S7.** The ESS {*F, a_g_ + a_p_*} when saved resources increase the reproductive resources in the next flowering season under stochasticity within flowering seasons. Unless specified, *m* = 0.05,, *F*_0_ = 50, *l* = 10, *β* = 1.2, *k_g_* = *k_p_* = 0.5, 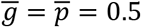, and *θ* = *θ_g_* = *θ_p_*, 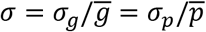. The initial strategy is *F* = 5, *a_g_ = a_p_* = 0.

| | $\bar{g} = \bar{p} = 0.2$ | $\bar{g} = \bar{p} = 0.5$ | $\bar{g} = \bar{p} = 0.8$ | $\bar{g} = 0.2,$<br>$\bar{p} = 0.8$ | $l = 5$ | $m = 0.2$ | $F_0 = 100$ |
| --- | --- | --- | --- | --- | --- | --- | --- |
| $\theta = 1, \sigma = 0.2$ | 8, 0.10 | 9, 0.14 | 10, 0.18 | 9, 0.14 | 24, 0.12 | 7, 0.18 | 20, 0.15 |
| $\theta = 0.01, \sigma = 0.2$ | 8, 0.10 | 9, 0.14 | 10, 0.18 | 9, 0.14 | 24, 0.12 | 7, 0.18 | 20, 0.15 |
| $\theta = 1, \sigma = 0.5$ | 8, 0.11 | 10, 0.15 | 10, 0.18 | 9, 0.14 | 24, 0.13 | 7, 0.18 | 19, 0.15 |
| $\theta = 0.01, \sigma = 0.5$ | 8, 0.11 | 10, 0.15 | 10, 0.18 | 9, 0.14 | 23, 0.12 | 7, 0.18 | 19, 0.15 |
| $\theta = 1, \sigma = 0.8$ | 8, 0.11 | 10, 0.16 | 10, 0.19 | 9, 0.15 | 24, 0.14 | 7, 0.19 | 19, 0.16 |
| $\theta = 0.01, \sigma = 0.8$ | 8, 0.11 | 10, 0.16 | 10, 0.19 | 9, 0.15 | 23, 0.12 | 7, 0.19 | 18, 0.15 |

**Table S8.**
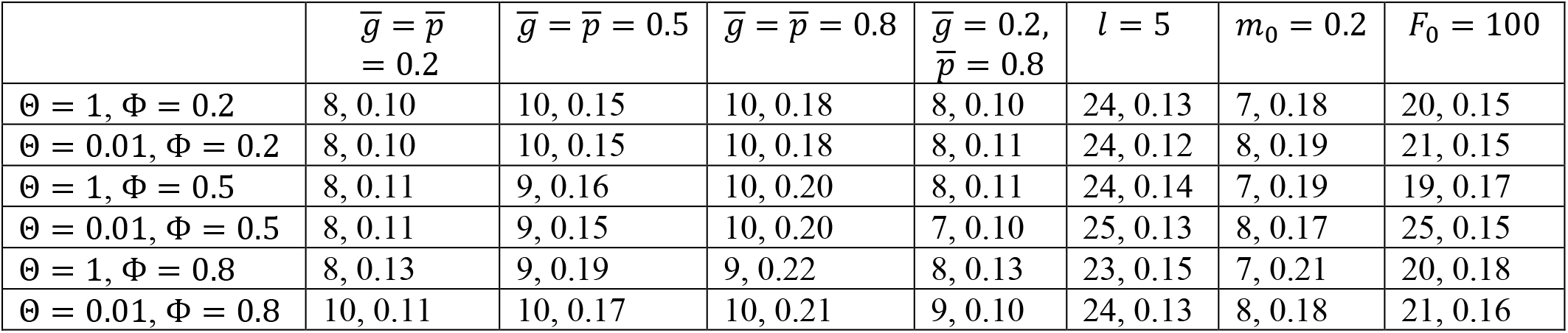
The ESS {*F, a_g_ + a_p_*} when saved resources increase the reproductive resources in the next flowering season under stochasticity between flowering seasons. Unless specified, *m* = 0.05, *F*_0_ = 50, *l* = 10, *β* = 1.2, *k_g_* = *k_p_* = 0.5, 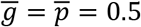, and Θ = Θ*_g_* = Θ*_p_*, 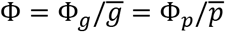. The initial strategy is *F* = 5, *a_g_* = *a_p_* = 0.

| | $\bar{g} = \bar{p} = 0.2$ | $\bar{g} = \bar{p} = 0.5$ | $\bar{g} = \bar{p} = 0.8$ | $\bar{g} = 0.2, \bar{p} = 0.8$ | $l = 5$ | $m_0 = 0.2$ | $F_0 = 100$ |
| --- | --- | --- | --- | --- | --- | --- | --- |
| $\Theta = 1, \Phi = 0.2$ | 8, 0.10 | 10, 0.15 | 10, 0.18 | 8, 0.10 | 24, 0.13 | 7, 0.18 | 20, 0.15 |
| $\Theta = 0.01, \Phi = 0.2$ | 8, 0.10 | 10, 0.15 | 10, 0.18 | 8, 0.11 | 24, 0.12 | 8, 0.19 | 21, 0.15 |
| $\Theta = 1, \Phi = 0.5$ | 8, 0.11 | 9, 0.16 | 10, 0.20 | 8, 0.11 | 24, 0.14 | 7, 0.19 | 19, 0.17 |
| $\Theta = 0.01, \Phi = 0.5$ | 8, 0.11 | 9, 0.15 | 10, 0.20 | 7, 0.10 | 25, 0.13 | 8, 0.17 | 25, 0.15 |
| $\Theta = 1, \Phi = 0.8$ | 8, 0.13 | 9, 0.19 | 9, 0.22 | 8, 0.13 | 23, 0.15 | 7, 0.21 | 20, 0.18 |
| $\Theta = 0.01, \Phi = 0.8$ | 10, 0.11 | 10, 0.17 | 10, 0.21 | 9, 0.10 | 24, 0.13 | 8, 0.18 | 21, 0.16 |

**Table S9.** The ESS {*F, a_g_ + a_p_*} when saved resources increase the reproductive resources in the next flowering season under stochasticity both within and between flowering seasons. Unless specified, *m* = 0.05, *F*_0_ = 50, *l* = 10, *β* = 1.2, *k_g_* = *k_p_* = 0.5, 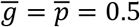. Also, *θ* = *θ_g_* = *θ_p_*, 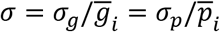, and Θ = Θ*_g_* = Θ*_p_*, 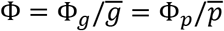. The initial strategy is *F* = 5, *a_g_* = *a_p_* = 0.

| | $\bar{g} = \bar{p} = 0.2$ | $\bar{g} = \bar{p} = 0.5$ | $\bar{g} = \bar{p} = 0.8$ | $\bar{g} = 0.2, \bar{p} = 0.8$ |
| --- | --- | --- | --- | --- |
| $\Theta = \theta = 1, \Phi = \sigma = 0.2$ | 9, 0.14 | 10, 0.15 | 10, 0.19 | 9, 0.15 |
| $\Theta = \theta = 0.01, \Phi = \sigma = 0.2$ | 9, 0.12 | 10, 0.15 | 10, 0.19 | 9, 0.14 |
| $\Theta = \theta = 1, \Phi = \sigma = 0.5$ | 9, 0.14 | 10, 0.19 | 10, 0.19 | 9, 0.17 |
| $\Theta = \theta = 0.01, \Phi = \sigma = 0.5$ | 9, 0.14 | 11, 0.15 | 13, 0.17 | 9, 0.15 |
| $\Theta = \theta = 1, \Phi = \sigma = 0.8$ | 9, 0.17 | 10, 0.20 | 9, 0.23 | 9, 0.21 |
| $\Theta = \theta = 0.01, \Phi = \sigma = 0.8$ | 9, 0.13 | 9, 0.16 | 10, 0.21 | 9, 0.16 |

**Table S10.**
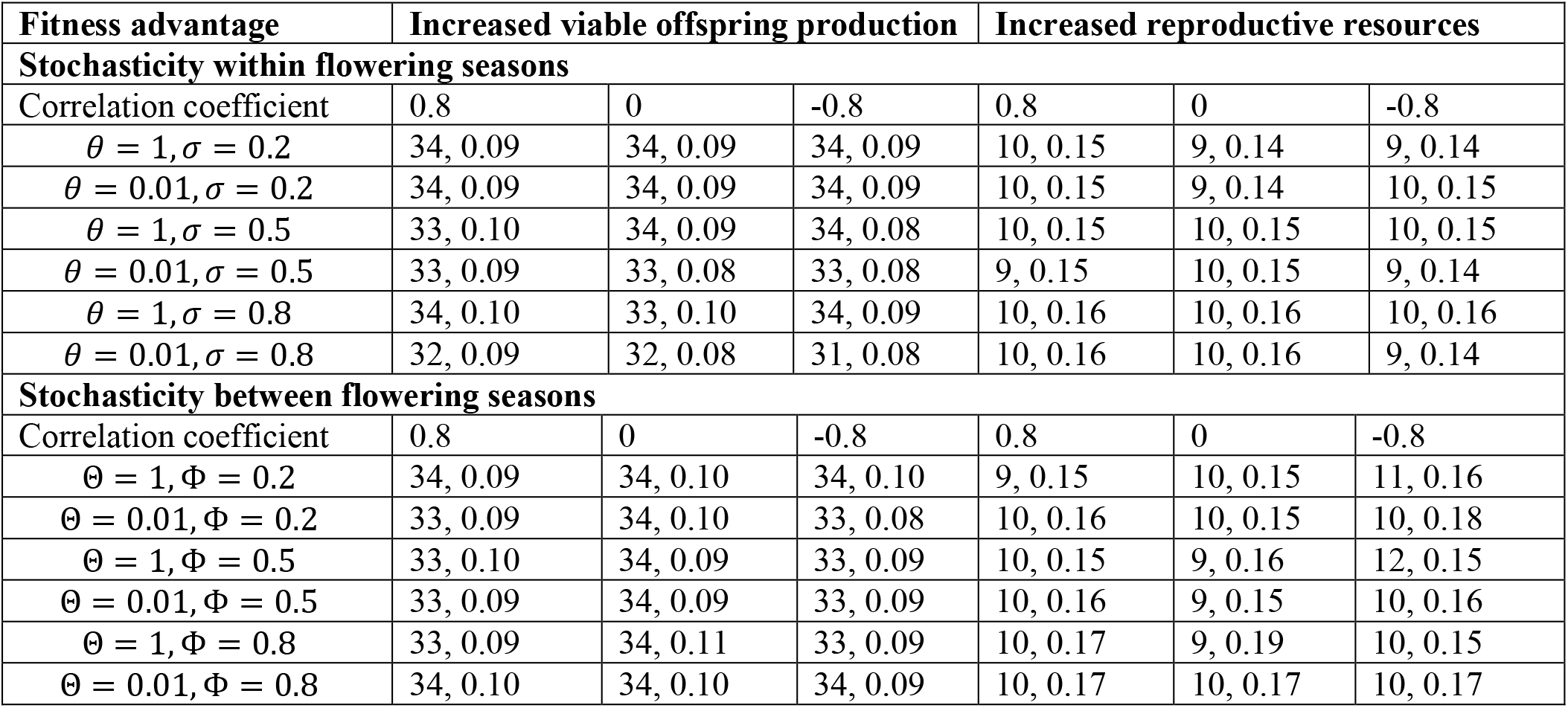
Effects of the correlation coefficient between male and female fitness accrual rates on the ESS {*F, a_g_* + *a_p_*} under stochasticity within or between flowering seasons. The initial population starts with the strategy *F* = 5, *a_g_* = *a_p_* = 0. For increased viable offspring production, *α* = 2, and for increased reproductive resources in the next flowering season, *β* = 1.2, and 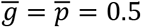, *k_g_ = k_p_* = 0.5.

